# Trade of commercial potting substrates: A largely overlooked means of the long-distance dispersal of plants

**DOI:** 10.1101/2021.12.01.470741

**Authors:** Judit Sonkoly, Attila Takács, Attila Molnár V., Péter Török

**Affiliations:** Department of Ecology, University of Debrecen, H-4032 Debrecen, Egyetem tér 1., Hungary; MTA-DE Lendület Functional and Restoration Ecology Research Group, H-4032 Debrecen, Egyetem tér 1., Hungary; Department of Botany, University of Debrecen, H-4032 Debrecen, Egyetem tér 1., Hungary

**Keywords:** cattle manure, dispersal, growing media, horticulture, human-vectored dispersal, invasions, long-distance dispersal, potting soil, potting substrate, seed dispersal, traits

## Abstract

Although long-distance dispersal (LDD) events are considered to be rare and highly stochastic, they are disproportionately important and drive several large-scale ecological processes. The realisation of the disproportionate importance of LDD has led to an upsurge in studies of this phenomenon; yet, we still have a very limited understanding of its frequency, extent and consequences. Humanity intentionally spreads a high number of species, but a less obvious issue is that it is associated with the accidental dispersal of other plant species. Although the global trade of potted plants and horticultural substrates is capable of dispersing large quantities of propagules, this issue has hardly been studied from an ecological point of view. We used the seedling emergence method to assess the viable seed content of different types of commercial potting substrates to answer the following questions: (i) In what richness and density do substrates contain viable seeds? (ii) Does the composition of substrates influence their viable seed content? and (iii) Are there common characteristics of the species dispersed this way? We detected altogether 438 seedlings of 66 taxa and found that 1 litre of potting substrate contains an average of 13.27 seeds of 6.24 species, so an average 20-litre bag of substrate contains 265 viable seeds. There was a high variability in the seed content of the substrates, as substrates containing cattle manure contained a substantially higher number of species and seeds than substrates without manure. Based on this, this pathway of LDD is an interplay between endozoochory by grazing livestock and accidental human-vectored dispersal, implying that the diet preference of grazing animals largely determines the ability of a plant species to be dispersed this way. According to our results, potting substrates can disperse large quantities of seeds of a wide range of plant species, moreover, these dispersal events occur on very long distances in almost all cases. We conclude that this kind of human-vectored LDD may have complex effects on plant populations and communities; however, as this dispersal pathway is largely understudied and has hardly been considered as a type of LDD, its consequences are still largely unknown and further studies of the issue are of great importance.

## Introduction

Understanding plant dispersal is becoming increasingly important in the face of the steadily increasing anthropogenic influence on natural habitats also including climate change (Thuiller et al. 2008, Renton et al. 2013). Climate change imposes increasing pressure on plant species to adjust their distribution range (Parmesan & Yohe 2003; Walther et al. 2005), while they also need to ensure their persistence in rapidly changing landscapes (Bossuyt & Honnay 2006). As a consequence, effective long-distance dispersal is likely more important for plant species than ever before.

Although long-distance dispersal (LDD) events are considered to be rare and highly stochastic, they are disproportionately important and drive several large-scale ecological processes (Nathan 2006, Auffret & Cousins 2013). Range shifts tracking a changing landscape and climate, the long-term persistence of species in fragmented habitats, gene flow between populations, and the spread of invasive species are among the several large-scale processes of great conservation concern mainly determined by LDD rather than local dispersal (Nathan 2006). Estimating local dispersal capacity and distances is challenging enough, not to mention quantifying LDD events (Levin et al. 2003). The realisation of the disproportionate importance of LDD has led to an upsurge in studies of this phenomenon; yet, we still have only a very limited understanding of its frequency, extent and consequences (Jordano 2017). The range of phenomena involved in LDD is not yet explored either (Higgins et al. 2003).

Humanity has been an important dispersal vector for plants throughout history with its importance increasing especially fast in recent times (Auffret 2011), making humanity one of the major (or perhaps the major) dispersal vector (Hodkinson & Thompson 1997). Agriculture and horticulture are among the main means of human-vectored dispersal (Hodkinson & Thompson 1997, Murray 2012), and ornamental plants are considered as the main source of alien plants, many of them becoming naturalized or even invasive in their new range (Haeuser et al. 2017, Hulme et al. 2018). A less obvious issue is that the increasing worldwide trade of cultivated plants and horticultural substrates (often referred to as growing media) is often associated with the dispersal of other contaminant species (Prach et al. 1995, Hodkinson & Thompson 1997).

There are sporadic studies reporting the occurrence of adventive species in nurseries (e.g., Hoste et al. 2009, Gallego & Lumbreras 2013), but the propagule content of container plant substrates and bagged commercial potting substrates has hardly been considered from an ecological point of view. Conn et al. (2008) studied the seed content of soils obtained from container-grown ornamentals in Alaska and identified 54 species, including several species invasive in Alaska. By analysing the viable seed content of commercial potting soils in the USA, Dyer et al. (2017) detected the seeds of 80 species and concluded that it would be necessary to reduce the presence of viable seeds in commercial potting soils in order to prevent further dispersal of weedy species. Composted cotton gin waste (or cotton gin trash), a by-product of cotton processing, is also used as a component of potting substrates in cotton-producing areas of the USA (Jackson et al. 2005). Norsworthy et al. (2009) demonstrated that a high number of seeds from several weed species can be found in cotton gin waste even after 2 years of composting; thus, using it as soil amendment or potting substrate constituent provides an effective pathway for the dispersal of seeds.

Given that horticultural substrates (or growing media) are produced in huge quantities (more than 30 million m^3^ produced every year within the EU, Schmilewski 2009) and constituent materials or even ready-to-use substrates are regularly imported from distant countries (Schmilewski 2009), the importance of this dispersal pathway is probably largely underestimated. As the propagule content of potting substrates and the associated potential LDD of plants represent a conspicuous knowledge gap, further investigations of this issue are critically important. We addressed this gap by setting up a seedling emergence experiment in which we assessed the viable seed content of different types of commercial potting substrates to answer the following questions: (i) In what richness and density do substrates contain viable seeds? (ii) Does the composition of substrates influence their viable seed content? and (iii) Are there common characteristics of the species dispersed this way?

## Materials & Methods

We purchased 3 bags of each of 11 different commercial potting substrates (for types and characteristics of the substrates see Table 1). We aimed to obtain potting substrates from several producer companies, as available on the market at the time. We took samples of 1-liter volume from each bag of potting substrate (33 l altogether). From here on, we handled the samples as if they were collected samples for analyzing the soil seed bank.

**Table 1.**
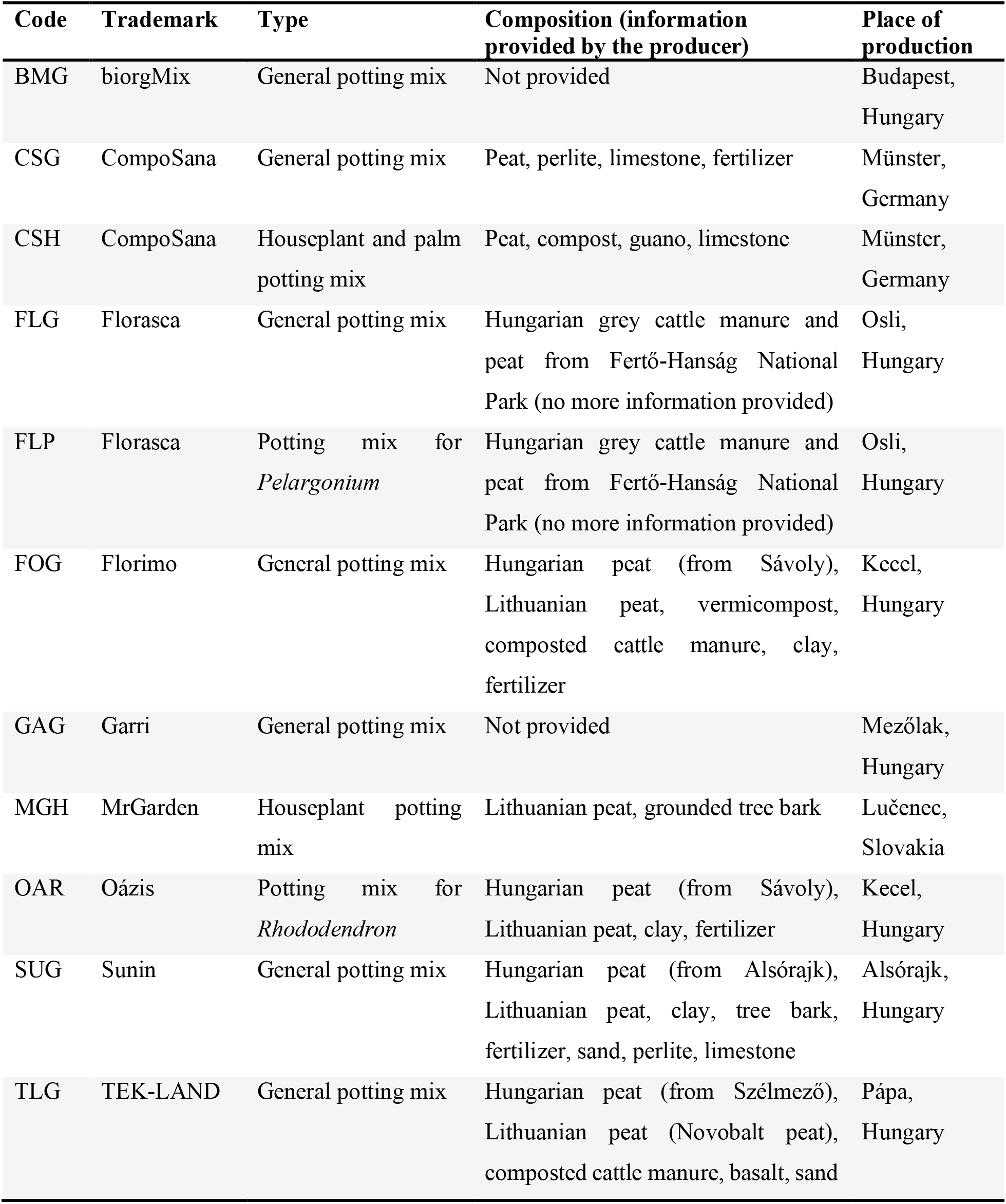
Types and characteristics of the studied potting substrates.

To reduce sample volume, we followed the sample concentration method of Ter Heerdt et al. (1996). Samples were washed through two sieves, this way washing out fine mineral and organic particles. The coarse mesh (2.8 mm) retained larger particles, while the fine mesh (0.2 mm) retained all the seeds and other particles of similar size. We filled 33 balcony planters (60 cm × 20 cm × 15 cm) with steam-sterilised potting substrate and spread a concentrated sample in a thin layer (up to 4-5 mm in depth) on the surface of each. Samples were kept in an unheated greenhouse from 22 March 2019 until 12 November 2019, and they were watered daily to provide optimal conditions for germination. Seedlings were identified, counted, and removed regularly, and the plants not yet identifiable were transplanted into separate pots (altogether 133 transplanted individuals) and grown until they developed flowers or other diagnostic features. We also set up five balcony planters as control, filled only with steam-sterilised potting substrate to detect possible air-borne seed contamination.

We also calculated the greatest distance each species dispersed in the substrates measured as a straight line between the place of substrate production and the place of purchase (Debrecen, Hungary). Then we compared this dispersal distance to the recorded or predicted maximum dispersal distance (MDD) of each species (Supplementary Material 1). Recorded MDDs were gathered from the literature by Tamme et al. (2014), while predicted MDDs were either calculated by Tamme et al. (2014) or calculated by us using the dispeRsal function in R (Tamme et al. 2014).

To compare the seed weight of the dispersed species and the regional flora, we obtained thousand-seed weight (TSW) data for the regional flora from published sources containing data of measurements carried out in Hungary (Schermann 1967, Csontos et al. 2003, 2007, Török et al. 2013, Török et al. 2016) (Supplementary Material 1).

Seed bank type data for the regional flora were also gathered from studies carried out in Hungary (Csontos 2001, Török 2008, Valkó et al. 2011, Tóth 2015, and Csontos et al. 2016) to avoid the confounding effect of different climatic and environmental conditions. Using the Seed Longevity Index (SLI) of Bekker et al. (1998), we express the ratio of data indicating a persistent soil seed bank with a value between 0 and 1, where 0 means that all available data indicate a transient seed bank and 1 means that all available data indicate persistent seed bank (Supplementary Material 1).

Leaf trait data, namely leaf area (LA), specific leaf area (SLA) and leaf dry matter content (LDMC) were obtained from published sources containing data of measurements carried out in Hungary (Lhotsky et al. 2016, E-Vojtkó et al. 2020), and leaf trait data of the species germinated from the samples were also supplemented with our own unpublished data (Supplementary Material 1).

To compare ecological indicator values for the dispersed species and the regional flora, we gathered the Borhidi-type indicator values of the regional flora for soil moisture (WB), light intensity (LB) and nutrient supply (NB) (Supplementary Material 1). Borhidi-type indicator values (Borhidi 1995) are based on Ellenberg indicator values (Ellenberg et al. 1992) but adapted to the Pannonian Ecoregion. Mann-Whitney U tests were used to compare the seed number and species number of substrate samples containing versus not containing cattle manure. Substrates for which the presence or absence of manure was not specified were not considered in the comparison. TSW, LA, SLA, LDMC, SLI, WB, LB and NB values of the dispersed species and the regional flora were also compared using Mann-Whitney U tests. All statistical analyses were conducted in R version 4.0.3 (R Core Team 2020).

## Results

Altogether 438 seedlings germinated from the samples of the commercial potting substrates, which belonged to 66 taxa (Supplementary Material 2). In some cases, plants could not be identified at the species level because they died or did not develop diagnostic features even after two vegetation seasons, but from here on we will use the term species for simplicity. The most abundant species in the samples were widespread, common species: *Portulaca oleracea, Capsella bursa-pastoris, Echinochloa crus-galli, Juncus effusus* and *Poa annua* (with 40, 34, 32, 28 and 28 seedlings, respectively).

More than half of the species (41 out of the 66 species) only occurred in one type of potting substrate or in different substrates produced by the same manufacturer (FLG and FLP, 16 of the species were only found in these two substrates, Supplementary Material 2, for substrate codes see Table 1). On the other hand, some of the species were particularly widespread among different substrates: *Juncus articulatus* and *Veronica catenata* were both found in 6 different substrates while *Echinochloa crus-galli, Plantago major* and *Portulaca oleracea* occurred in 5 substrates.

The mean number of seeds in 1 liter of substrate was 13.27 (min=0, max=61). At least one seed germinated from each type of substrate, but not from each 1-liter sample. The mean number of species in 1 liter of substrate was 6.24 (min=0, max=21) (Table 2). There was a substantial difference between the seed content of different substrates (Table 2). The highest seed density and species diversity was observed in the substrates FLG and FLP produced by the same manufacturer (Fig. 1). Samples of these two substrates contained 270 seeds of 39 species, accounting for more than half of all the seeds and more than half of all the species.

**Table 2.** The total number of seedlings and species in the samples of each type of potting substrate, and the mean number of seedlings and species in 1 liter of each substrate. For the coding of the substrates see Table1.

| Code | Total seedling nr. | Seedlings per liter | Total species nr. | Species per liter |
| --- | --- | --- | --- | --- |
| BMG | 31 | 10.33 | 17 | 7.33 |
| CSG | 2 | 0.67 | 1 | 0.33 |
| CSH | 1 | 0.33 | 1 | 0.33 |
| FLG | 108 | 36.00 | 27 | 16.00 |
| FLP | 162 | 54.00 | 31 | 18.00 |
| FOG | 6 | 2.00 | 4 | 2.00 |
| GAG | 30 | 10.00 | 18 | 7.33 |
| MGH | 4 | 1.33 | 3 | 1.00 |
| OAR | 2 | 0.67 | 2 | 0.67 |
| SUG | 36 | 12.00 | 14 | 7.33 |
| TLG | 56 | 18.67 | 16 | 8.33 |

**Fig. 1.**
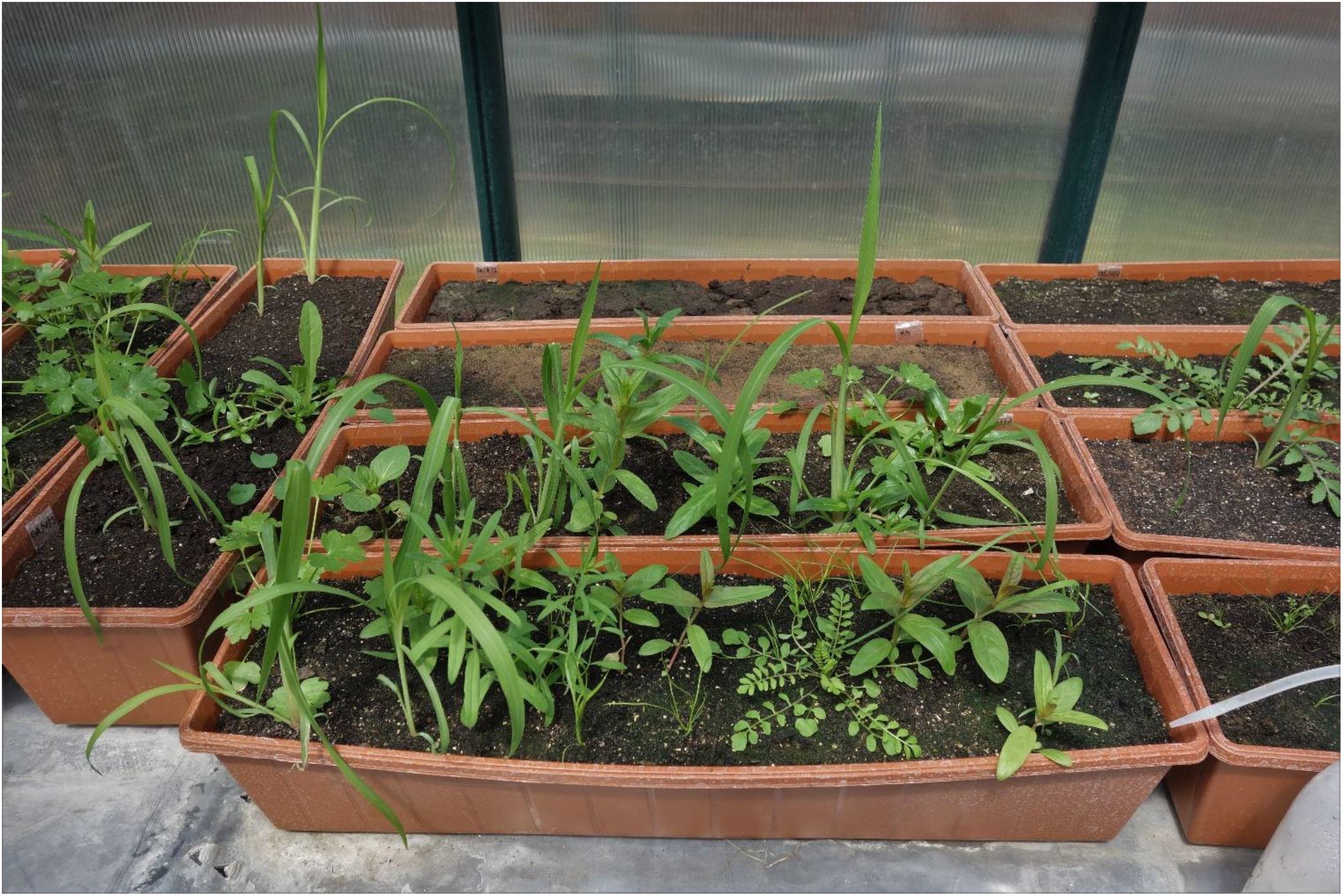
A part of the germination experiment with a sample of the potting substrate (FLG) with the highest seed and species number in the front.

Substrates containing cattle manure had a substantially higher seed number (Mann-Whitney U test, W=18, p<0.001) and species number (Mann-Whitney U test, W=17, p<0.001) than substrates not containing manure (Fig. 2).

**Fig. 2.**
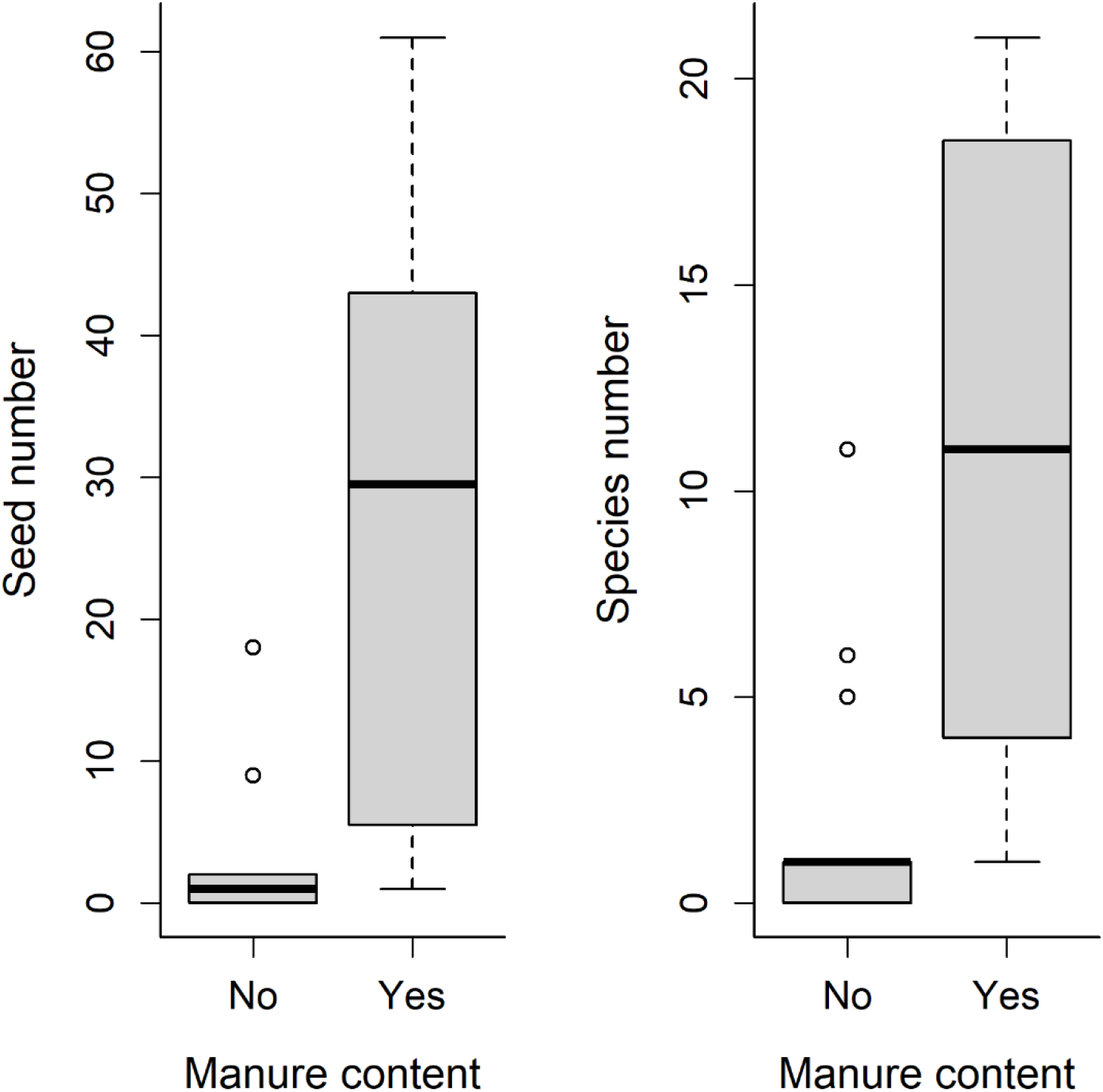
Seed number and species number in 1-liter samples of potting substrates containing versus not containing cattle manure.

The actual distances the seeds dispersed in the potting substrates from the place of production to the place where the substrates were purchased ranged from 171 km (Lučenec, Slovakia) to 1117 km (Münster, Germany) (Fig. 3). On average, seeds travelled 311.7 km in the substrates. The distances the species dispersed in the substrates were, on average, more than 300,000 times greater (min. 28×, max. 2,243,527×) than the previously recorded or predicted maximum dispersal distances for these species (Supplementary Material 1). However, it should be considered that the distances dispersed by these seeds may be even greater as some constituents of the substrates originate at locations even farther (see Table 1), but in this case we have no way of knowing which seed came from which constituent material. Another thing to consider is that these substrates are presumably transported and sold at locations even farther from their place of production.

**Fig. 3.**
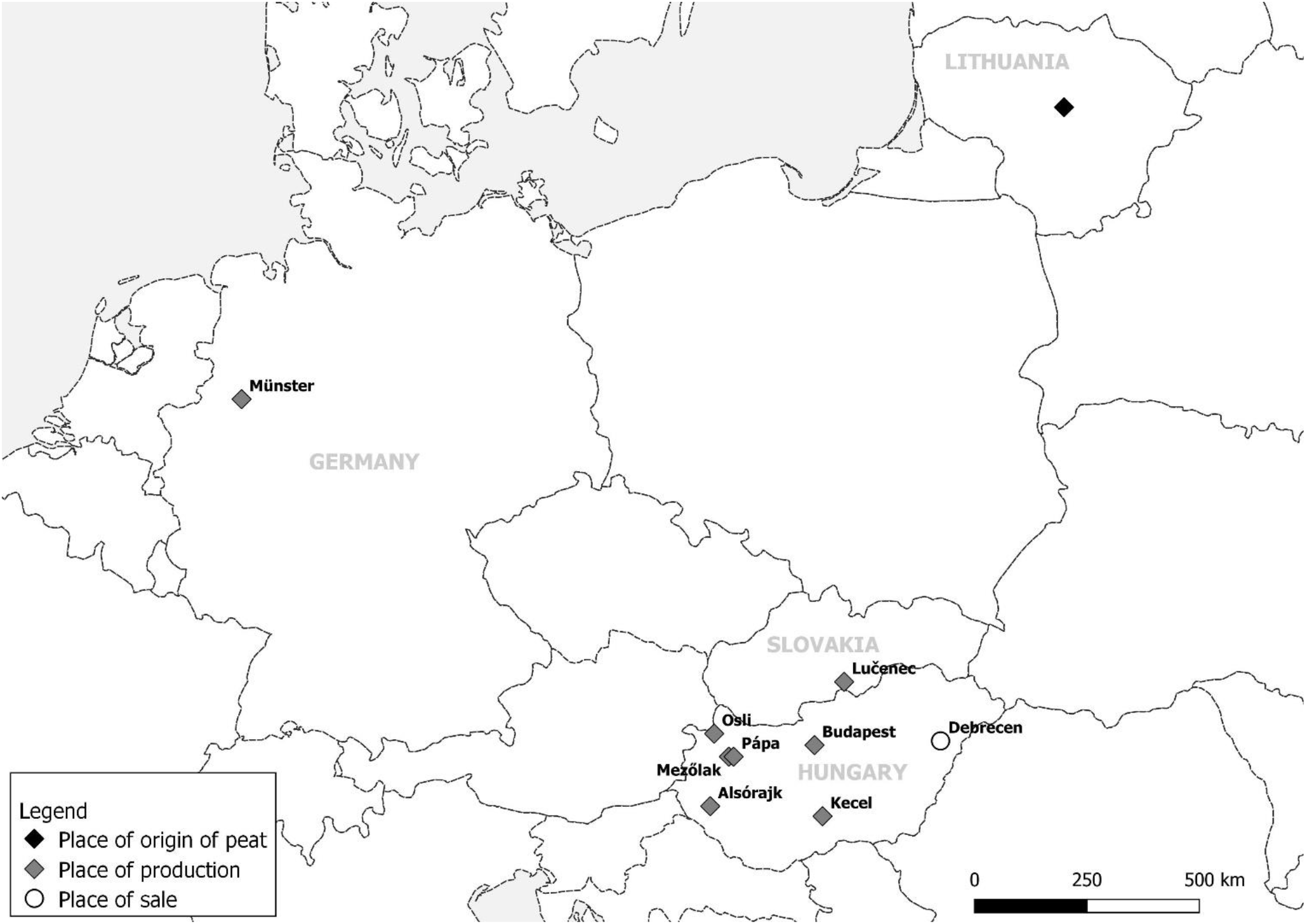
Map showing the studied potting substrates’ place of production, the place where they were purchased, and Lithuania, the country of origin of peat used in several types of potting substrates.

The dispersed species did not differ significantly from the regional flora in terms of leaf area, leaf dry matter content and light intensity indicator values, but did differ in terms of thousand-seed weight, seed longevity index, specific leaf area, and indicator values for soil moisture and nutrient supply (Table 3). In other words, species detected in the potting substrates generally had smaller and more persistent seeds, and higher specific leaf area than the regional flora, and were characterized by higher indicator values for soil moisture and nutrient supply.

**Table 3.**
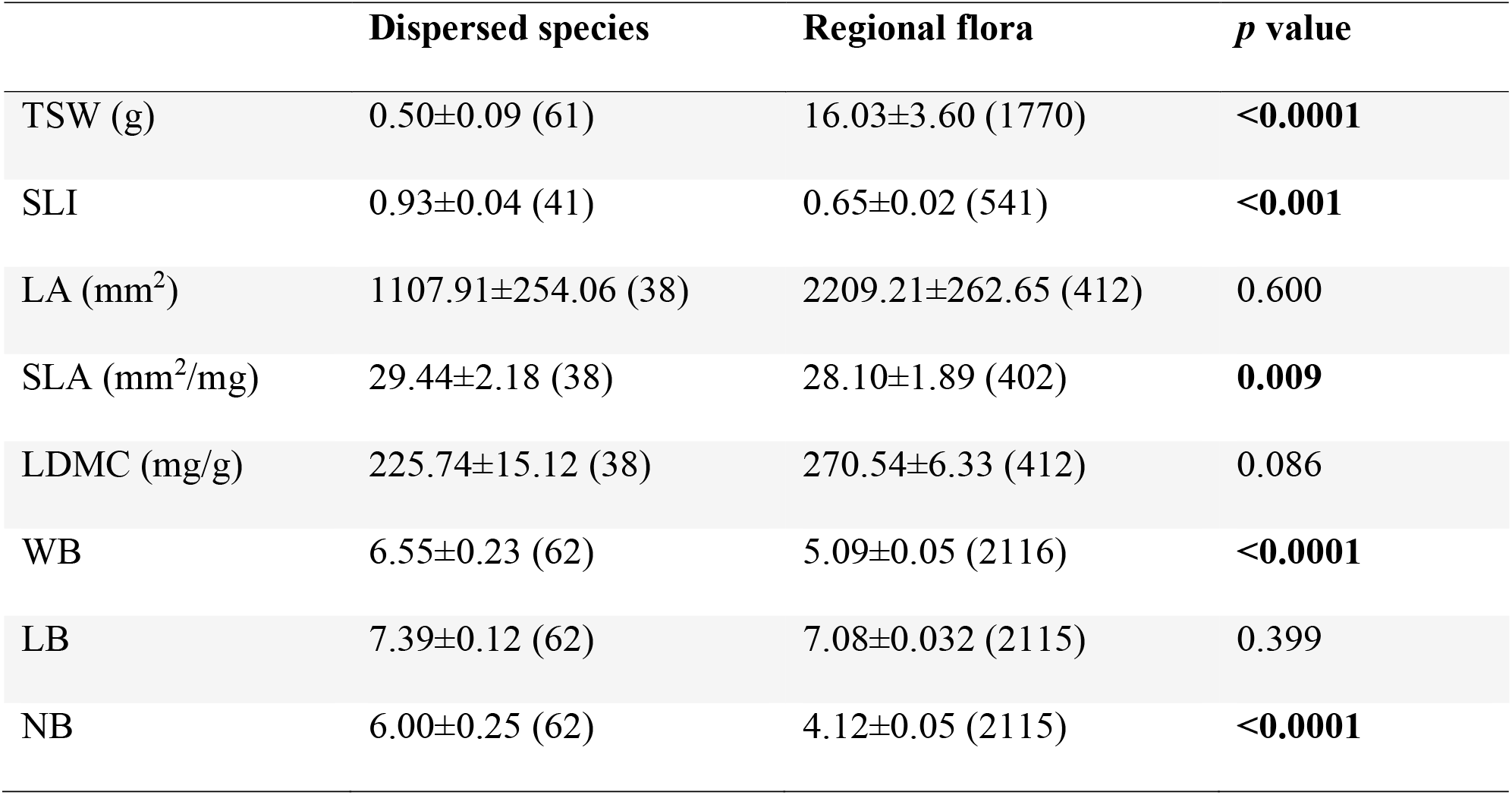
Thousand-seed weight (TSW), seed longevity index (SLI), leaf area (LA), specific leaf area (SLA), leaf dry matter content (LDMC), and Borhidi-type ecological indicator values for soil moisture (WB), light intensity (LB) and nutrient supply (NB) compared between the species detected in the potting substrates and species of the regional flora. Mean±SE (sample size), Mann-Whitney U tests, significant differences are indicated by boldface.

## Discussion

Although horticultural substrates have a great potential to disperse plant propagules, this pathway of long-distance dispersal (LDD) has been largely overlooked so far (Hodkinson & Thompson 1997). Despite their potential importance in the dispersal of plants (and also animals, see Moore et al. 2016), the viable seed content of commercial potting substrates remained strikingly understudied so far (Dyer et al. 2017). Our results show that a regular 20-l bag of potting substrate contains an average of 265.4 viable seeds (min: 6.6 seeds/20 l; max: 1080 seeds/20 l). Given the annual production of more than 30 million m^3^ of horticultural substrates within 13 countries of the EU alone (Schmilewski 2009), substrates produced within the EU may disperse an estimated 398 billion of seeds annually. This number is certainly just a rough estimate, but it highlights the order of magnitude of seeds dispersed this way and confirms that this is not a negligible phenomenon.

To provide a more accurate estimation of the number of seeds dispersed by potting substrates, further factors, first of all the composition of the substrates should be considered. In line with the increasing awareness of environmental issues, the use of composted organic waste (e.g., composted plant material and manure) in potting substrates is expected to rise to replace non-renewable constituents like peat (Ceglie et al. 2015, Barret et al. 2016). Given that our results suggest that manure content is the most important factor influencing the viable seed content of potting substrates, increasing use of manure is expected to further increase the potential of substrates to contain and disperse seeds. Other organic (e.g., peat, coconut coir, tree bark, and wood fibre) and inorganic materials (mostly clay and perlite) commonly used in substrate production (Schmilewski 2009) most probably contain very low amounts of viable seeds. A more accurate estimation is also hindered by the fact that the volume of global horticultural substrate production is not known, but probably increasing.

As the majority of seeds are dispersed on short distances (up to few dozen meters, e.g., Howe & Smallwood 1982, Cain et al. 2000), rare events of seeds traveling on long distances are disproportionately important and influence key ecological processes (Cain et al. 2000, Nathan 2006). What constitutes long-distance dispersal (LDD) is up to debate and depends on the context. It can be defined based on a small proportion of seeds that were dispersed on the longest distances, or a pre-defined threshold distance can be set as an absolute definition (Nathan et al. 2008), ideally based on an ecologically meaningful distance, such as the typical distance between distinct populations or suitable habitat patches (Higgins et al. 2003). For example, Cain et al. (2000) considered dispersal events over 100 m as LDD, while Higgins & Richardson (1999) considered seeds travelling 1–10 km as LDD in their simulations. Based on this, seed dispersal via potting substrates constitutes LDD in virtually every case, so the importance of the high number of seeds dispersed this way is especially high. The effectiveness of this dispersal pathway is further increased by the conditions at the destination often being ideal for germination and establishment (Hodkinson & Thompson 1997, Conn et al. 2008), such as in the substrate of an ornamental plant transplanted to a garden, or potting substrates ending up in gardens or even in nature (Foxcroft et al. 2008, Rusterholz et al. 2012)

Five of the species that germinated from the substrate samples were non-native in Hungary, although four of them are already particularly widespread in Hungary (*Amaranthus retroflexus, Conyza canadensis, Erigeron annuus* and *Solidago gigantea*), and only *Epilobium ciliatum* is still rather sporadic in the country, while several native, but sporadic species were found in the samples (e.g., *Cardamine parviflora, Hypericum tetrapterum, Juncus subnodulosus, Rumex maritimus, Thalictrum flavum* and *Veronica catenata*; see Bartha et al. 2021). Based on this, besides potentially introducing alien species to the local flora, potting substrates presumably have an important role in transporting the seeds of native species between distant populations and into new, previously unoccupied habitats. Therefore, seeds dispersed by these substrates may play an important role in gene flow between distant populations and range shifts tracking a changing landscape and climate (Travis et al. 2013).

An important aspect of our findings is that manure content is the most significant factor determining the ability of potting substrates to contain and disperse seeds, implying that dispersal by horticultural substrates constitutes an interplay of endozoochory and accidental human-vectored dispersal (Bullock & Pufal 2020). Consequently, the diet preference of grazing animals, i.e., how likely a plant species is to be consumed by grazers (mostly cattle), largely determines the ability of a plant species to be dispersed this way (Cosyns et al. 2005, Picard et al. 2016). Our finding that species detected in the substrates had higher specific leaf area compared to the regional flora can also be explained by the foraging strategy of grazing animals, which usually prefer plant species with high specific leaf area due to the higher nutritional value and palatability of such species (Pontes et al. 2007, Lloyd et al. 2010). The ability of seeds to survive passing through the digestive tract of animals is also a critical factor, which is confirmed by the fact that species that germinated from the substrate samples typically had small, persistent seeds, which traits generally promote gut passage and endozoochorous dispersal (e.g., Albert et al. 2015, Picard et al. 2016). As most of the species we detected in the substrates do not have specialized structures for long-distance dispersal (e.g., special structures for anemochorous or zoochorous dispersal, see Correira et al. 2018 or Chen et al. 2020), our findings support the notion that dispersal syndromes are inadequate for predicting plant species’ ability for LDD (Green et al. 2021).

In landscapes heavily modified by humans, the loss of some key dispersal services, such as the loss of large wild herbivores, herded livestock and large frugivores (e.g., Mokany et al. 2014, Bakker et al. 2016, Tucker et al. 2018) induces a loss of LDD events which is a serious conservation issue (Jordano 2017). On the other hand, in such heavily modified landscapes, increased human activity can take over the role of standard dispersal vectors, and humans are already considered as the most important LDD vectors (Higgins et al. 2003, Nathan 2006). Globalization, specifically the steeply rising rate of global trade and human mobility has led to the increased dispersal of a diverse array of organisms over large distances (e.g., Westphal et al. 2008, Hulme et al. 2009), implying that increasing human activity and globalization resulted in increased chances of LDD. Here we showed that, besides potentially introducing new alien species, the unintentional dispersal of seeds and other propagules by the increasing global horticultural trade probably constitutes a more general LDD pathway for a large number of species. As such, horticultural trade can affect other large-scale ecological processes of great conservation concern, such as gene flow between otherwise isolated populations and range shifts tracking the changing climate and landscape. Thus, this kind of human-vectored LDD may have complex effects on plant populations and communities, including both adverse and favourable consequences. However, as this dispersal pathway is largely understudied and has hardly been considered as a type of LDD, its consequences are still largely unknown and further studies of the issue are of great importance.

## Acknowledgements

We would like to thank T. Abonyi, N. Balogh, K. Süveges and E. Tóth-Szabó for their help during the experiment. JS was supported by NKFIH-OTKA PD 137747, AT and AMV were supported by NKFIH-OTKA K132573, and PT was supported by NKFIH-OTKA K119225 and K137573.

## Supplementary Materials

Supplementary Material 1: Characteristics of species detected in the potting substrates.

Supplementary Material 2: The number of seeds of each species found in 1-L samples.

